# Exploring the archaeome: detection of archaeal signatures in the human body

**DOI:** 10.1101/334748

**Authors:** Manuela R. Pausan, Cintia Csorba, Georg Singer, Holger Till, Veronika Schöpf, Elisabeth Santigli, Barbara Klug, Christoph Högenauer, Marcus Blohs, Christine Moissl-Eichinger

## Abstract

Due to their fundamentally different biology, archaea are consistently overlooked in conventional microbiome surveys. Using amplicon sequencing, we evaluated methodological set-ups to detect archaea in samples from five different body sites: respiratory tract (nose), digestive tract (mouth, appendix, and stool) and skin. With the optimized protocols, the detection of archaeal ribosomal sequence variants (RSVs) was increased from one (found in currently used, so-called “universal” approach) to 81 RSVs in a representative sample set. In order to assess the archaeome diversity, a specific archaea-targeting methodology is required, for which we propose a standard procedure. This methodology might not only prove useful for analyzing the human archaeome in more detail but could also be used for other holobionts’ samples.

## Introduction

The importance of microbial communities to human and environmental health motivates microbiome research to uncover their diversity and function. While the era of metagenomics and metatranscriptomics has begun, 16S rRNA gene amplicon sequencing still remains one of the most used methods to explore microbial communities, mainly due to the relatively low cost, the number of available pipelines for data analysis, and the comparably low computational power required.

It has been recognized that methodological issues in sample processing can significantly influence the outcome of microbiome studies, affecting comparability between different studies ^1,2^ or leading to an over-and under-estimation of certain microbial clades ^3,4^. For better comparability among different studies, standard operational procedures for sampling, storing samples, DNA extraction, amplification and analysis were set-up (e.g. the Earth Microbiome Project ^5^ and the Human Microbiome Project ^6^). This includes the usage of so-called “universal primers” ^7–9^, to maximally cover the broadest prokaryotic diversity.

The human microbiome consists of bacteria, archaea, eukaryotes and viruses. The overwhelming majority of microbiome studies is bacteria-centric, but in recent years, awareness on eukaryotes (in particular fungi) and viruses has increased ^10–12^. However, most microbiome studies still remain blind for the human archaeome ^3,13^. A few of the underlying reasons for the under-representation of archaea in microbiome studies are (i) primer mismatches of the “universal primers” ^14^, (ii) the sometimes too low abundance of the archaeal DNA in the studied samples ^15^, (iii) improper DNA extraction methods ^16^, and (iv) the incompleteness of the 16S rRNA gene reference databases due to missing isolates, especially for the DPANN superphylum ^15,17^. Moreover, the clinical interest on archaea is minor, due to the fact that there are no known or proved archaeal pathogens yet ^18^.

Nevertheless, (methanogenic) archaea are part of the commensal microorganisms inhabiting the human body, being regularly detected in the oral cavity and the gastrointestinal tract ^19–22^; in the latter they sometimes even outnumber the most abundant bacterial species (14%, ^23^). Most human archaea studies use either cultivation or qPCR methods ^24–30^ and only a few, 16S rRNA gene sequencing archaea-centric studies are available ^24,31–33^. These new studies have shown that archaea are also present in the human respiratory tract ^24^ and on human skin in considerable amounts ^31,34^. Furthermore, Koskinen et al. ^24^ have shown for the first time that archaea reveal a body site specific pattern, similar to bacteria: the gastrointestinal tract being dominated by methanogens, the skin by *Thaumarchaeota*, the lungs by *Woesearchaeota*, and the nose archaeal communities being composed of mainly methanogens and *Thaumarchaeota*. Altogether, this indicates a substantial presence of archaea in some, or even all, human tissues.

As a logic consequence of our previous studies, we have started to optimize the detection a methods of archaea as human commensals. We tested, *in silico* and experimentally, 27 different 16S rRNA gene targeting primer pair combinations suitable for NGS amplicon sequencing, to detect the archaeal diversity in samples from different body sites, including respiratory tract (nose samples), digestive tract (oral samples, appendix specimens and stool), and skin. Our results culminate in a proposed standard operating procedure for archaea diversity analysis in human samples.

## Results

Primer pairs were evaluated with respect to the following characteristics: high *in silico* specificity for archaeal 16S rRNA genes and an amplicon length of 150 to 300 bp, suitable for NGS, and *in vitro* capability to amplify diverse archaeal 16S rRNA genes from a variety of human specimens

Besides archaea-specific primer pairs, two widely used “universal” primers (515F-806uR original; 515FB-806RB modified; ^7,9^) were evaluated all along to assess the potential of “universal” primers to display archaeal diversity associated with the human body.

### Most archaea-targeting primers reveal good coverage in silico

A total of 12 different primer pairs were evaluated *in silico* (Table 1). Most primer pairs showed high coverage for the archaeal domain ranging from 46% to 89% and revealed a high domain-specificity (8 of 12 primer pairs without matches outside of the archaeal domain). When one mismatch per primer was allowed, the coverage increased to values from 68% to 95%.

**Table 1.**
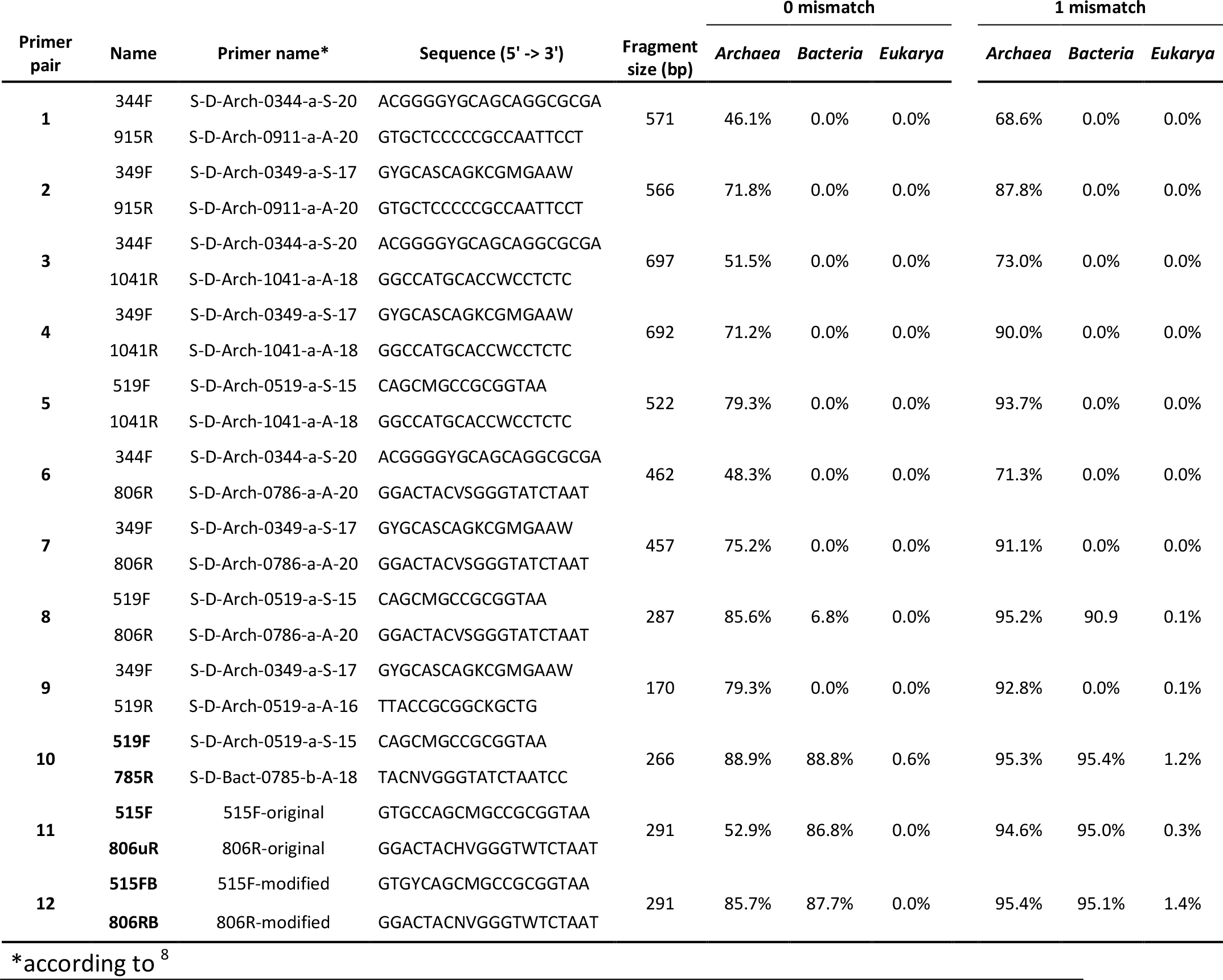
Primer selection and results of the pre-analysis *in silico* evaluation of all primer pairs used. Coverage of *Archaea*, *Bacteria* and *Eukarya* is given in percentages, depending on whether no or one mismatch was allowed. Designated “universal” primers (primer pairs 10-12) are indicated in bold letters.

One designated archaeal primer pair was found to target additionally sequences of the bacterial and eukaryotic domain when one mismatch per primer was allowed, namely primer pair 519F-806R, with a coverage of the bacterial domain > 90%.

We further evaluated the detailed coverage of the primer pairs for specific archaeal phyla and genera of particular interest in human archaeome studies: *Euryarchaeota*, *Thaumarchaeota*, and *Woesearchaeota*, as well as *Nitrososphaera, Methanobrevibacter*, *Methanosphaera* and *Methanomassiliicoccus*. For all subsequent *in silico* analyses we allowed one mismatch.

All primer pairs revealed a high coverage for the *Euryarchaeota* phylum (in total >90%), for genera *Methanobrevibacter* (between 94.6% and 98.9%) and *Methanomassiliicoccus* (between 92.9% and 100%), while the coverage for *Methanosphaera* was below 90% for most primer pairs except for 519F-806R and 349F-519R (Table 2).

**Table 2:** *In silico* analysis of the coverage of chosen primer pairs for specific archaeal taxa of interest. One mismatch was allowed per primer. For primer full names and sequences, please refer to Table 1.

| primer pair | Name | *Euryarchaeota* |  |  |  | *Thaumarchaeota* |  | *Nanoarchaeota* |
| --- | --- | --- | --- | --- | --- | --- | --- | --- |
|  |  | total | *Methano-brevibacter* | *Methano-sphaera* | *Methano-massiliicoccus* | total | *Nitrososphaera* | (*Woesearchaeota*) |
| 1 | 344F |  |  |  |  |  |  |  |
|  | 915R | 90.80% | 95.30% | 82.20% | 100.00% | 20.60% | 87.60% | 66.40% |
| 2 | 349F |  |  |  |  |  |  |  |
|  | 915R | 91.50% | 95.30% | 84.20% | 100% | 92% | 89.70% | 70.30% |
| 3 | 344F |  |  |  |  |  |  |  |
|  | 1041R | 90.80% | 94.60% | 79.40% | 100% | 20.70% | 89.00% | 67.90% |
| 4 | 349F |  |  |  |  |  |  |  |
|  | 1041R | 91.50% | 94.60% | 79.40% | 100% | 96.40% | 92.30% | 74.30% |
| 5 | 519F |  |  |  |  |  |  |  |
|  | 1041R | 95% | 97.80% | 85.40% | 92.90% | 96.60% | 90.60% | 83% |
| 6 | 344F |  |  |  |  |  |  |  |
|  | 806R | 92.30% | 95.50% | 82.60% | 100% | 23.30% | 88% | 65.20% |
| 7 | 349F |  |  |  |  |  |  |  |
|  | 806R | 93.20% | 95.60% | 84.20% | 100% | 96.50% | 90.10% | 72.90% |
| 8 | 519F |  |  |  |  |  |  |  |
|  | 806R | 96.60% | 98.90% | 90% | 95% | 96.70% | 89.50% | 83.10% |
| 9 | 349F |  |  |  |  |  |  |  |
|  | 519R | 93.60% | 95.80% | 90.70% | 95% | 98% | 94.40% | 83.10% |
| 10 | 519F |  |  |  |  |  |  |  |
|  | 785R | 96.50% | 98.60% | 89.60% | 95% | 96.20% | 87.80% | 87.60% |
| 11 | 515F |  |  |  |  |  |  |  |
|  | 806R | 96.20% | 98.60% | 89.60% | 95% | 94.70% | 86.90% | 89.50% |
| 12 | 515FB |  |  |  |  |  |  |  |
|  | 806RB | 96.20% | 98.60% | 89.60% | 95% | 96.50% | 89% | 89.50% |

The coverage of the *Thaumarchaeota* phylum depended on the primer pair used. Most analyses that included the primer 344F showed a low *in silico* coverage for *Thaumarchaeota* (below 30%) while all other primer pair combinations revealed a high coverage of this phylum (>90%; Table 2). The coverage for *Nitrososphaera* in particular varied between 86.9% and 94.4%. The class *Woesearchaeota* showed variable coverage between 65.2% and 89.5%.

As the archaeal primer 344F has often been used for detecting archaea in a variety of environmental samples ^35,36^, we took a closer look on its coverage capacity using the TestProbe 3.0 ^8^ and the SILVA database SSU132 ^37^. Overall, the primer revealed 73.2% coverage of the archaeal domain. The *in silico* results showed a high coverage of the *Euryarchaeota* phylum (93.8%) and the genera within, especially *Methanobrevibacter* with 96.1%, *Methanosphaera* with 89.9% and *Methanomassiliicoccus* with 100%. It also revealed a good coverage for *Woesearchaeota* with 74.6%, but showed, despite a high coverage for the genus *Nitrososphaera* (93.6%), a generally low coverage of the *Thaumarchaeota* phylum with only 24%, indicating a potentially low capacity for studies with thaumarchaeotal diversity in focus.

Another primer that we analyzed in more detail was primer 519F, also known as S-D-Arch-0519-a-S-15. As the sequence of this primer (5’ - CAGCMGCCGCGGTAA - 3’) overlaps with the sequence of the “universal” primer S-*-Univ-0519-a-S-18 (5’ - CAGCMGCCGCGGTAATWC - 3’), we were interested to compare their coverages.

As expected, the results from the *in silico* analysis indicated that the primer S-D-Arch-0519-a-S-15 targets *Bacteria* (coverage 98%), *Archaea* (coverage 98.2%) and *Eukarya* (coverage 96.4%). The universal primer S-*-Univ-0519-a-S-18 has a similar coverage and specificity for the three domains of life: *Bacteria* (coverage 97.5%), *Archaea* (coverage 96.4%), and *Eukarya* (coverage 95.6%). Considering our *in silico* results, the primer S-D-Arch-0519-a-S-15 cannot be used to target archaea specifically and should be re-named to S-D-Univ-0519-a-S-15.

As most selected archaea-targeting primers revealed a good coverage of the archaeal domain in general, all primer pairs were used for subsequent wet-lab experiments.

### Archaeal community composition varies according to the used primer pairs and universal primers fail to detect the archaeal diversity

Herein we sought to identify the optimal primer pair for amplicon sequencing of the archaeomes in human samples. For this, we selected five representative sample types from different body sites: nose (upper nasal cavity), oral (subgingival sites), stool and appendix specimens, and skin (back) (sample set 1). The stool sample represented the natural positive control.

Next generation sequencing was performed, after a two-step nested PCR (for archaea) or a single-step PCR (“universal” target). The nested PCR approach was selected based on the reasons given in the Materials and Methods section. In brief, the first PCR was intended to select the archaeal community of interest, the second to further amplify the archaeal signal.

The use of universal primers (primer pair 515F-806uR, 515FB-806RB and 519F-785R) in the PCR reaction resulted in reads that were classified mainly within the bacterial domain with almost no reads classified within the archaea, confirming our previous observations ^24^. In fact, when the two universal primer pairs (515F-806uR original and 515FB-806RB) were compared regarding the archaeal domain, only primer pair 515F-806uR allowed the detection of only one RSV being classified within the archaea and from only one sample, the stool sample.

Universal primer pair 519F-785R yielded slightly better results, allowing the detection of three different archaeal RSVs from two different samples: *Methanobrevibacter* and *Methanosphaera* in the stool sample, and one RSV from the nose sample, classified within the *Thaumarchaeota* phylum. Very similar results (detection of the same methanoarchaeal signatures in the stool sample, and one thaumarchaeal signature in the oral sample instead of the nose sample) were obtained from primer pair 519F-806R, which was originally described to be archaea-specific, but revealed wide coverage of the bacterial and archaeal domain (>90%, when one mismatch allowed) *in silico* (see previous chapter).

To identify whether the universal primer pairs allow the detection of the same RSVs or closely related RSVs in the analyzed samples, a phylogenetic tree was constructed (Fig. 1). Besides the obtained archaeal RSVs from the universal approaches, the RSVs retrieved from the archaeal specific primer pair combination 344F-1041R/519F-806R were included for comparison. This approach allowed the detection of 20 RSVs in the nose, 19 RSVs in the oral, one RSV in the appendix, 3 RSVs in the stool, and 39 RSVs in the skin sample. For the stool sample, the RSVs obtained from the universal and archaeal specific approach grouped together, either within *Methanobrevibacter* or *Methanosphaera* clade (Fig. 1), whereas the RSVs (universal and specific approach) from nose and oral samples diversified.

**Fig. 1:**
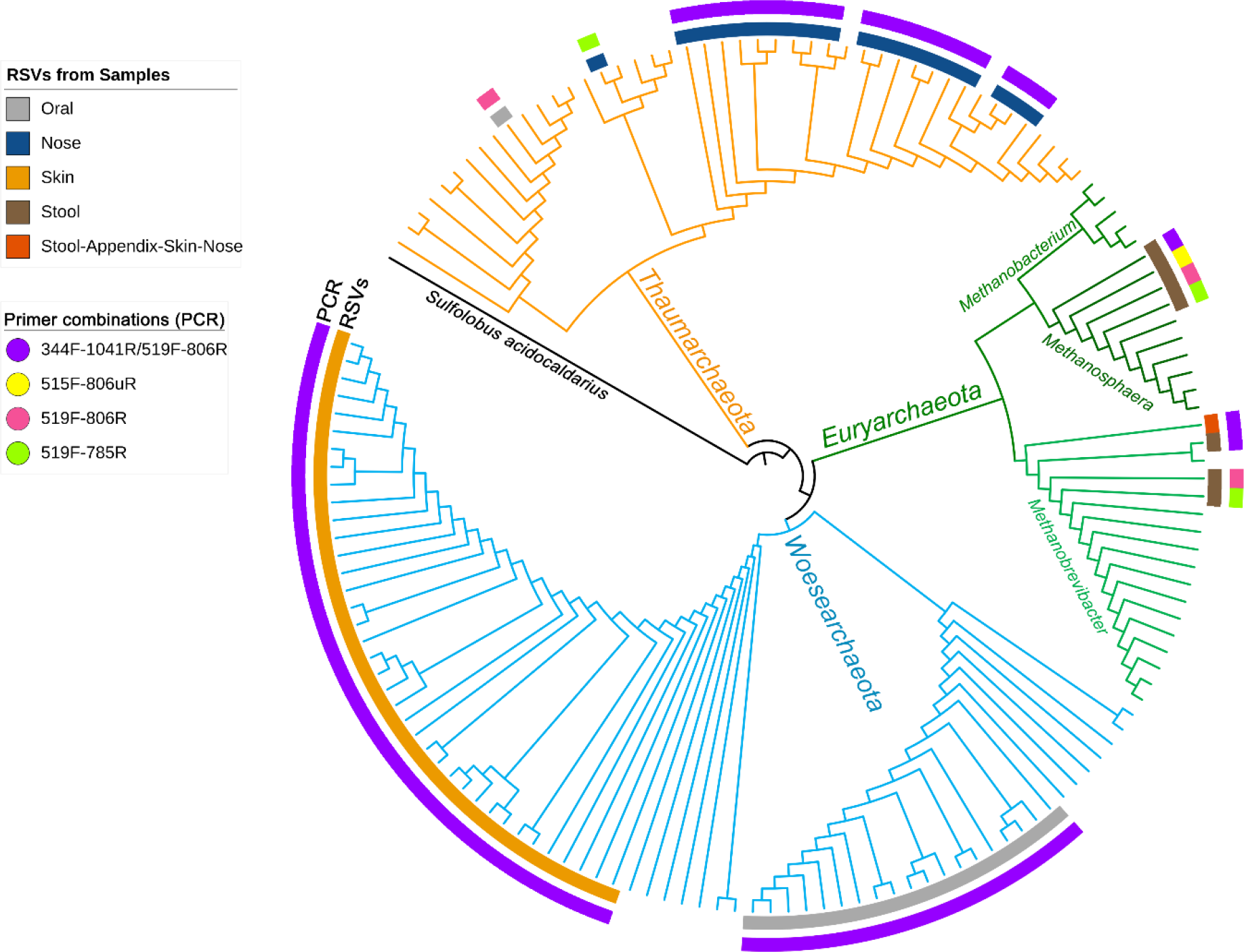
Phylogenetic tree based on the retrieved RSVs from the universal approach, archaeal approach with primer 519F-806R or from the PCR based on the primer pair combination 344F-1041R/519F-806R as indicated in colors as an outermost circle (legend “Primer combinations (PCR)”). The inner circle represents the body site from where the RSVs were identified (see legend). Reference sequences from the SILVA database are shown without label. The branches of the tree were colored according to the phyla, blue: *Woesearchaeota*, green: *Euryarchaeota*, and orange: *Thaumarcheota*.

Overall, 10 out of 23 primer pair combinations allowed the detection of archaeal signatures in all analyzed samples. All 23 primer pair combinations were able to detect archaeal reads in at least one of the sample types analyzed, for example all primer pair combinations detected archaeal RSVs in the stool sample; the number of RSVs, however, varied according to the used primer pair combination.

Depending on the used primer pair, the archaeal community composition was found to be highly variable (Suppl. Fig. 1). We observed that the detected variation in the archaeal composition was due to the used primer pair in the first PCR, the primer pair used to select the communities, while the second PCR and primer pair enhanced the signal of the first PCR (Suppl. Fig. 1). It shall be mentioned that for the second PCR only three different primer pairs have been used, 349F-519R, 519F-785R and the 519F-806R, of which the first two primer pairs had been used before to explore archaeal communities in human samples ^24^ and in confined habitats ^39^.

To further explore the influence of the primer pair selection on the archaeal community composition, the alpha diversity was calculated using the Shannon index (Fig. 2). For this analysis, we excluded the results obtained from the second primer pair 349F-519R as most samples herein (except stool samples) yielded less than 500 reads.

**Fig. 2:**
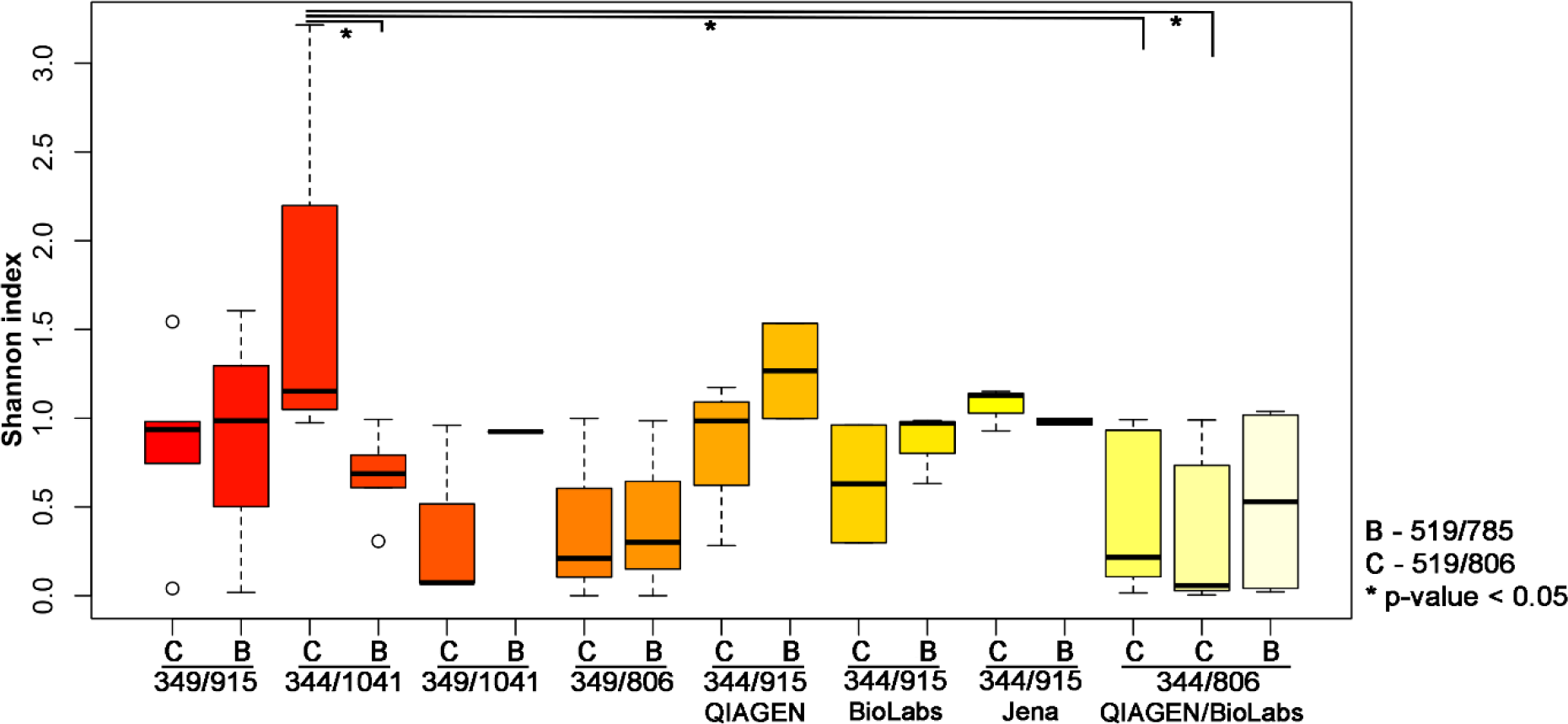
Shannon index indicating the diversity received from different PCR approaches. The results have been plotted and grouped according to the first PCR used and the statistical significance (p-value <0.05; Wilcoxon Rank Test) is indicated by *.

The highest archaeal diversity could be detected with the primer combination 344F-1041R/519F-806R (PCR34); this result was found to be significant (p<0.05) compared to PCR 33 (344F-1041R/519F-785R), PCR Q7 (344F-806R/519F-806R) and PCR M7 (344F-806R/519F-806R; Table 3 and Fig. 2), whereas no other significant differences could be detected.

**Table 3.** displays all primer pair combinations used for the first and the second PCR of the nested approach and the “universal” PCR. If not indicated otherwise (in brackets), the first PCR was followed by a purification of the PCR product by the MinElute PCR Purification kit (QIAGEN) kit. n.a.: not applicable.

| PCR # | Primer combination<br>1st PCR | Primer combination<br>2nd PCR |
| --- | --- | --- |
| PCR21 | 349F-915R | Illu 349F-Illu519R |
| PCR22 |  | Illu 519F-Illu785R |
| PCR23 |  | Illu 519F-Illu806R |
| PCR31 | 344F-1041R | Illu 349F-Illu519R |
| PCR33 |  | Illu 519F-Illu785R |
| PCR34 |  | Illu 519F-Illu806R |
| PCR41 | 349F-1041R | Illu 349F-Illu519R |
| PCR42 |  | Illu 519F-Illu785R |
| PCR43 |  | Illu 519F-Illu806R |
| PCR61 | 349F-806R | Illu 349F-Illu519R |
| PCR62 |  | Illu 519F-Illu785R |
| PCR63 |  | Illu 519F-Illu806R |
| PCR71 | 519F-1041R | Illu 519F-Illu785R |
| PCR72 |  | Illu 519F-Illu806R |
| PCR81 | 519F-806R | Illu 519F-Illu785R |
| PCR82 |  | Illu 519F-Illu806R |
| PCR91 | 344F-519R | Illu 349F-Illu519R |
| PCRQ1 | 344F-915R<br>(QIAGEN) | Illu 349F-Illu519R |
| PCRQ3 |  | Illu 519F-Illu785R |
| PCRQ4 |  | Illu 519F-Illu806R |
| PCRM1 | 344F-915R<br>(NEB Monarch) | Illu 349F-Illu519R |
| PCRM3 |  | Illu 519F-Illu785R |
| PCRM4 |  | Illu 519F-Illu806R |
| PCRA1 | 344F-915R<br>(Analytik Jena) | Illu 349F-Illu519R |
| PCRA3 |  | Illu 519F-Illu785R |
| PCRA4 |  | Illu 519F-Illu806R |
| PCRQ5 | 344F-806R<br>(QIAGEN) | Illu 349F-Illu519R |
| PCRQ6 |  | Illu 519F-Illu785R |
| PCRQ7 |  | Illu 519F-Illu806R |
| PCRM5 | 344F-806R<br>(NEB Monarch) | Illu 349F-Illu519R |
| PCRM6 |  | Illu 519F-Illu785R |
| PCRM7 |  | Illu 519F-Illu806R |
| PCR8-Uni | n.a. | Illu 515F-Illu806uR |
| PCR9-Uni |  | Illu 515FB-Illu806RB |
| PCR10 |  | Illu 519F-Illu806R |
| PCR11-Uni |  | Illu 519F-Illu785R |

According to the comparison of the alpha diversity of the archaeal communities between the different primer pair combinations, we recommend the use of the nested approach with the primer pair 344F-1041R in the first PCR, followed by a second PCR with the primers 519F-806R for studying and exploring the archaeal communities in human samples.

The use of the different purification kits between the first and the second PCR resulted in no significant results based on the alpha diversity (Shannon index) comparison using the Wilcoxon Rank Test (p-value >0.05; Fig. 2). Due to visible bands on the gel electrophoresis for the results obtained after the purification with the Monarch^®^ PCR & DNA Cleanup Kit (5 μg) (New England Biolabs GmbH; Ipswich, USA) we decided to further use this kit for the purification step.

### The primer combination with superior performance revealed a broad archaeal diversity in stool, appendix, nose, oral and skin samples

To further test and validate the use of the primer pair combination 344F-1041R/519F-806R for studying the archaeal communities within human samples, we selected additional samples from the same body sites: nose (n=5), oral (n=6), appendix (n=5), stool (n=5), and skin (n=7) (sample set 2).

Our selected PCR approach allowed the detection of archaea in all samples investigated with an average of 102,366 reads and 8 observed RSVs for the nose, 56,480 reads and 35 observed RSVs for oral, 46,022 reads and 8 observed RSVs for the appendix, 93,948 reads and 4 observed RSVs for the stool sample, and 76,001 reads and 30 observed RSVs for the skin samples.

The results were plotted to indicate the archaeal communities present at genus level in the analyzed samples (Fig. 3).

**Fig. 3:**
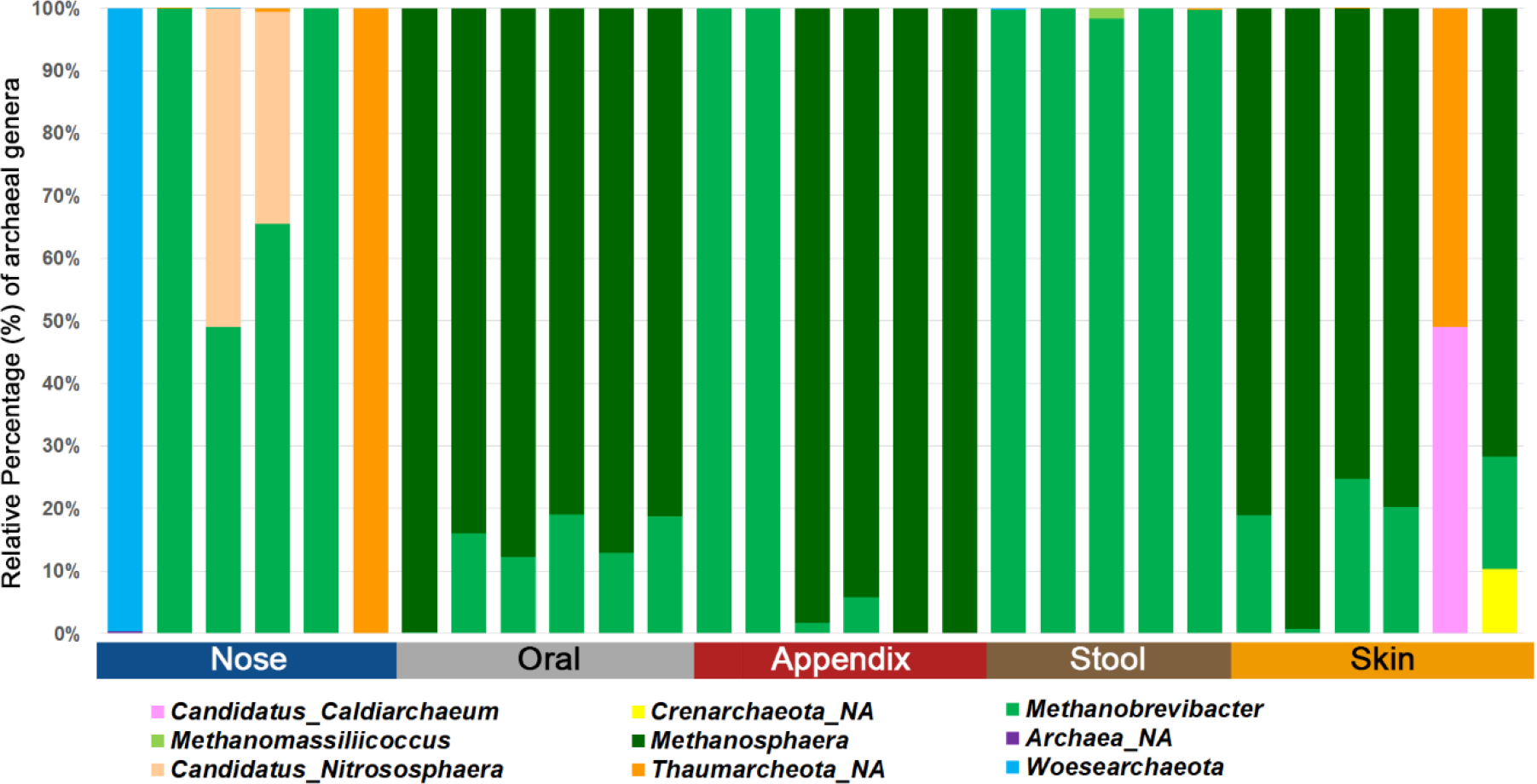
Bar chart displaying the different archaeal genera detected in different human samples using the superiorly performing primer combination 344F-1041R/519F-806R.

We further characterized the archaeal community information with respect to alpha and beta diversity. Depending on the body site a significant difference (p-value < 0.05) could be shown for alpha (Shannon index and richness) and beta diversity (PCoA and RDA) (Fig. 4). Our results confirm the findings that archaeal communities are body site specific ^24^.

**Fig. 4:**
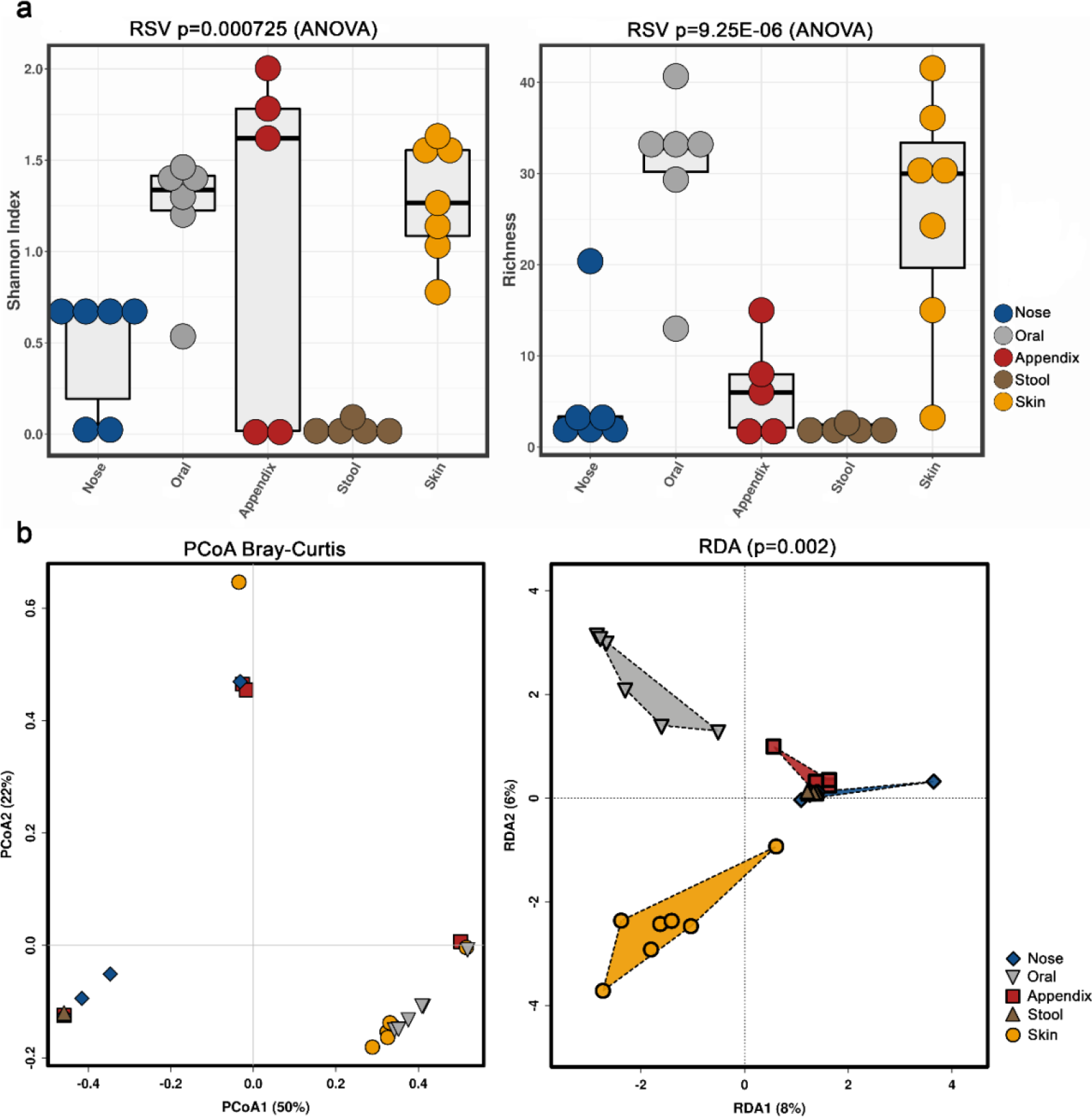
Alpha (a; Shannon index and richness) and beta diversity (b; PCoA and RDA) analyses of the obtained archaeal community information, based on primer combination 344F-1041R/519F-806R.

Notably, the stool samples revealed the overall lowest diversity of archaea, with only 3-5 identified archaeal RSVs, while skin and oral samples contained a higher diversity, with 5 to 49 RSVs found in the skin samples and 14 to 49 RSVs in the oral samples.

## Discussion

Up to now, little it is known about the composition of the human archaeome. It is unknown, whether archaeal communities are affected by dysbiosis or human disease, or how we acquire these microorganisms after birth, although several studies have shown that archaea are present in the first year of life ^27,40^. Additionally, it is largely unexplored, how archaeal communities interact/communicate with other commensal microorganisms inhabiting the human body. Furthermore, there still remains the most burning question, if there are really no archaeal pathogens. Facing these numerous unsolved mysteries, we argue that more studies are needed with respect to the human archaeome. For these, however, standardized protocols are required, which are powerful enough to reliably assess archaeal diversity and abundance based on 16S rRNA gene signatures.

To address the need for archaea-targeted amplicon method for NGS in human samples, we herein tested 12 different primers previously described in literature ^8^, in 27 primer pair combinations and evaluated their performance using *in silico* and experimental approaches on five different human sample types.

Despite their overall good *in silico* results, the three universal primer pairs tested failed to assess the archaeal diversity in the experiments. Two of these primer pairs represent the most-used universal primers for amplicon sequencing methods ^7,9^, resulting in the detection of one (515F-806uR) or zero archaeal RSVs (515FB-806RB) in five sample types that evidentially possessed a variety of archaeal signatures.

The reasons for the failure of the universal primers to detect Archaea are unclear; however, it seems bacterial signatures outcompete archaeal signatures, just due to slightly better primer matches, depending on the diversity within the sample.

Furthermore, an archaeal primer pair (519F-806R) that has been used before for amplicon sequencing ^41^ detected only a small proportion of the archaeal diversity in the analyzed samples, but the same primer pair performed better when used in a nested PCR together with the primer pair 344F-1041R for the first PCR.

Nested PCR has been shown to improve sensitivity and specificity and are useful for suboptimal DNA samples ^42,43^. Based on our experience in the past ^24^, other reports ^44^, and due to the fact that all attempts to use Illumina-tagged archaeal primers to directly identify archaeal 16S rRNA genes in human samples failed, we kept to this approach for the archaeal diversity assessment.

We used a combination of an archaea-specific first PCR (9 different primer combinations) and two archaeal specific and one universal primer pair, resulting in 23 different approaches (Table 3).

Notably, although the primer pair combinations 344F-915R/349F-519R and 344F-915R/519F-785R had been used earlier to detect archaeal signatures in human samples and confined environments ^24^ ^39^, our study revealed that when the second PCR contained the Illumina-tagged primers 349F-519R, almost no reads apart from the stool samples were retrieved.

Ten out of the 23 different primer combinations allowed the detection of archaeal signatures in all analyzed samples (sample set 1). The results of two of the primer pair combinations were outstanding regarding the number of reads and observed RSVs identified in each sample, namely primer pair 344F-1041R/519F-806R and 344F-1041R/519F-785R. The comparison of the alpha diversity (based on Shannon index) indicated that the archaeal diversity uncovered with the primer pair 344F-1041R/519F-806R was significantly higher than the one obtained with the primer pair combination 344F-1041R/519F-785R (Fig. 2), which was thus considered superior.

To further test and validate the use of the primer pair 344F-1041R/519F-806R, we selected 29 samples from different body sites (nose, oral, appendix, stool, skin; sample set 2), resulting in overall 85 archaeal RSVs from 6 different phyla. We were able to confirm body-site specificity through PCoA and RDA analysis ^24^, with the gastrointestinal tract (stool and appendix samples) being dominated by euryarchaeal communities, the oral samples dominated by archaeal communities from the *Euryarchaeota* phylum but different from the ones found in the gastrointestinal tract and the nose dominated by *Euryarchaeota* and *Thaumarchaeota* signatures. The skin revealed a mixture of *Euryarchaeota*, *Thaumarchaeota*, *Aenigmarchaeota*, and, in very low amounts also *Crenarchaeota*, confirming previous results ^24,31,45^.

According to the obtained results we recommend the use of the primer pair combination 344F-1041R/519F-806R to identify and characterize archaeal communities within human samples, even though the second primer pair 519F-806R is a universal primer pair according to the *in silico* results. Although this led to retrieval of not only archaeal reads, but also reads classified within *Bacteria* and *Eukarya* which had to be filtered bioinformatically, this procedure proved superior to all the other primer pairs tested in identifying archaeal signatures in the analyzed samples.

In conclusion, we have shown that the choice of the archaeal primer pair influences substantially the perspective of the obtained archaeal community in the analyzed samples. Therefore, for future comparisons between studies focused on exploring and characterizing the archaeal community in human samples using amplicon sequencing approach, it should be considered to make use of the same, standardized methodology. For this we recommend the use of a nested approach with the primer pair 344f-1041R for the first PCR, followed by a second PCR with the primer pair 519F-806R.

### Conclusions

The optimized and evaluated protocol for archaeal signature detection can now be used for all human samples and might also be useful for samples from other environments and holobionts, such plants or animals.

## Material and methods

### Selection of samples and DNA extraction

Representative sample types from various body sites including the respiratory tract (nose swabs), the digestive tract (oral biofilm, appendix biopsy and stool samples) and skin swabs were selected for the comparison of amplification-based protocols (See NOTE).

The nose swabs were obtained from healthy adults’ volunteers (18-40 years old) and were taken from the olfactory mucosa located at the ceiling of the nasal cavity using ultra minitip nylon flocked swabs (Copan, Brescia, Italy; n=7) ^46^. The oral samples have been obtained by standardized protocol for paper point sampling ^47^ from healthy children (10 years old) who participated in a microbiome study investigating the subgingival biofilm formation (n=7) ^48^. Appendix samples have been obtained during pediatric appendectomies from either acute or ulcerous appendicitis from children (7-12 years old) (n=6). Stool samples have been obtained from healthy adults’ volunteers (18-40 years old) (n=5) ^49^, and from one patient (68 years old) with above average methane production after metronidazole treatment (n=1; this sample was used for comparing different amplification protocols). Skin samples were obtained from healthy adults’ volunteers (18-40 years old) from either the back (n=1; this sample was used for comparing different amplification protocols) or the left forearm, using BD Culture Swabs^TM^ (Franklin Lakes, New Jersey, USA; n=7).

In all cases, the genomic DNA was extracted by a combination of mechanical and enzymatic lysis. However, depending on the sample type, different protocols were used: for the stool samples around 200mg of sample has been used for DNA extraction using the E.Z.N.A. stool DNA kit according to the manufacturer’s instruction. The DNA from the appendix samples was obtained using the AllPrep DNA/RNA/Protein Mini Kit (QIAGEN), before the DNA extraction, small pieces of cryotissue were homogenized 3 times for 30s at 6500rpm using the MagNALyzer ^®^ instrument (Roche Molecular Systems) with buffer RTL and β-mercaptoethanol (according to the manufacturer’s instructions). For the nose and skin samples from the forearm, the DNA was extracted using the FastDNA Spin Kit (MP Biomedicals, Germany) according to the provided instructions. The DNA from the oral samples and from the skin samples from the back were isolated using the MagnaPure LC DNA Isolation Kit III (Bacteria, Fungi; Roche, Mannheim, Germany) as described by Santigli et al. ^48^ and Klymiuk et al. ^50^.

**NOTE: Sample set 1** (one representative sample from each body site: nose, oral, appendix, stool from patient with high methane production and skin from the back) was used to initially evaluate the primers and methods, whereas **sample set 2** (6 nose samples, 6 oral samples, 5 appendices, 5 stool samples, and 7 skin samples) was then used for assessing the archaeal diversity, based on the chosen, optimized protocol.

### 16S rRNA gene primer selection and pre-analysis *in silico* evaluation

Different primer pairs targeting the archaeal 16S rRNA gene region have been selected from recent publications ^8,24^. The main criteria for selection were: a. specificity for archaea *in-silico*, b. low or no amplification of eukaryotic DNA, and c. amplicon length between 150 to 300bp, suitable for NGS such as Illumina MiSeq. In addition, three “universal” primer pairs ^7–9^ were tested in parallel to determine their efficiency in detecting archaea in human samples. Full information on the selected primer pairs is given in Table 1.

*In silico* evaluation of the selected primer pairs has been performed using the online tool TestPrime1.0 ^8^ and the non-redundant SILVA database SSU132 ^37^. Two of the primers (344F and S-D-Arch-0519-a-S-15) were also tested using TestProbe 3.0 ^8^ and the SILVA database SSU132 to assess their individual coverage for the archaeal domain. These two primers were tested either due to low coverage of the Thaumarchaeota domain (such as primer combinations including the 344F primer) or because the primers were targeting other domains of life such as Bacteria and Eukarya (primer combinations including the S-D-Arch-0519-a-S-15).

### PCR and library preparation

For archaea-targeting PCR, a nested approach was chosen to increase the specificity for archaea and to avoid the formation of primer dimers caused by the tag, necessary for Illumina sequencing, attached to the primers ^24,51^.

In addition to the nested approach, a standard PCR was performed with three different universal primer pairs, and one archaeal primer pair for comparative reasons, and to test if a universal approach is capable to cover archaea in human samples in sufficient depth. All primer combinations (in total 27) used for the PCR reactions are provided in Table 3.

For the first PCR, each reaction was performed in a final volume of 20 µl containing: TAKARA Ex Taq^®^ buffer with MgCl_2_ (10 X; Takara Bio Inc., Tokyo, Japan), primers 500 nM, BSA (Roche Lifescience, Basel, Switzerland) 1 mg/ml, dNTP mix 200 µM, TAKARA Ex Taq^®^ Polymerase 0.5 U, water (Lichrosolv^®^; Merck, Darmstadt, Germany), and DNA template (1-50 ng/µl).

After the first PCR, the resulting amplicons were purified to remove primer remnants. This purification was performed with three different kits to compare the different yields and efficiencies, namely MinElute PCR Purification kit (Qiagen; Hilden, Germany), Monarch^®^ PCR & DNA Cleanup Kit (5 μg) (New England Biolabs GmbH; Ipswich, USA), or innuPREP DOUBLEpure Kit (Analytik Jena, Germany) as indicated in Table 4. The purified PCR product was eluted in 10 µl water (Lichrosolv^®^; Merck, Darmstadt, Germany).

**Table 4:**
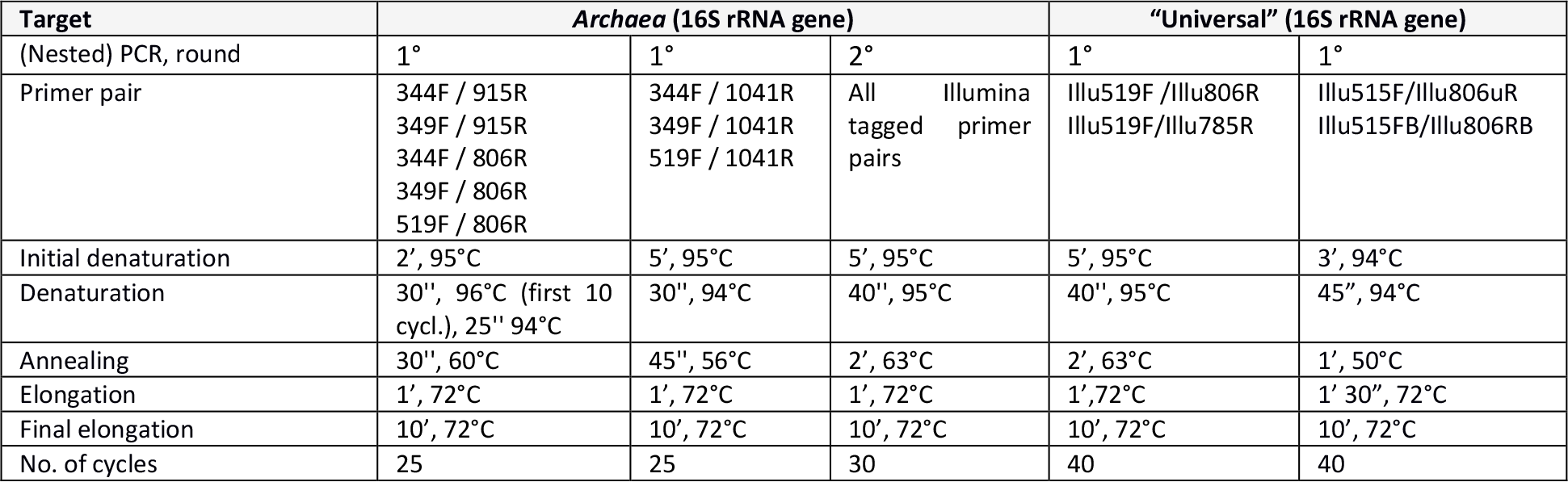
PCR conditions. For denaturation, annealing and elongation the corresponding time and temperature is given.

Two µl of the resulting, purified PCR products were transferred into a subsequent 2^nd^ PCR containing the following mixture: TAKARA Ex Taq^®^ buffer with MgCl_2_ (10 X; Takara Bio Inc., Tokyo, Japan), primers 500 nM, BSA (Roche Lifescience, Basel, Switzerland) 1 mg/ml, dNTP mix 200 µM, TAKARA Ex Taq^®^ Polymerase 0.5 U, and water (Lichrosolv^®^; Merck, Darmstadt, Germany) up to a volume of 25 µL.

The PCR cycling conditions are listed in Table 4, according to the primer pairs used. For all primer pairs, annealing temperatures were either determined experimentally by gradient PCR or adopted from literature information.

Sample set 2 was amplified using the primer combination 344F-1041R/519F-806R (Table 3). For the first PCR, each reaction was performed in a final volume of 20 µl as described above. After the first PCR, the PCR products were purified using Monarch^®^ PCR & DNA Cleanup Kit (5 μg; New England Biolabs GmbH). For the second PCR, the final volume was 25 µl, as described above, only the volume of the DNA template varied: 2 µl purified PCR product for stool and nose samples, 4 µl for all other samples.

### Next generation sequencing, bioinformatics and statistical analyses

Amplicons were sequenced at the ZMF Core Facility Molecular Biology in Graz, Austria, using the Illumina MiSeq platform ^50^. The MiSeq amplicon sequence data was deposited in the European Nucleotide Archive under the study accession number PRJEB27023.

The data processing of the obtained MiSeq sequence data was performed using the open source package DADA2 (Divisive Amplicon Denoising Algorithm; ^38^) as described previously ^39^. Shortly, the DADA2 turns paired-end fastq files into merged, denoised, chimera-free, and inferred sample sequences called ribosomal sequence variants (RSVs). The taxonomic affiliations were determined using SILVA v128 database as the reference database ^37^. In the resulting RSV table, each row corresponds to non-chimeric inferred sample sequence with a separate taxonomic classification.

Negative controls (extraction controls and no-template controls) were included during PCR amplification. The RSVs overlapping the negative controls and samples were either subtracted or completely removed from the data sets.

Processing of sequencing data was performed using the in-house Galaxy set-up ^50^ and subsequent statistical analyses were performed in R version 3.4.3 ^52^. Samples were rarefied to 500 reads and alpha diversity was calculated using the Shannon index. In order to identify differences between the archaeal diversity, Wilcoxon Rank Test was performed. The diversity of the archaeal communities within sample set 2 was determined using two diversity matrices (Shannon and richness). Analysis of variance (ANOVA) was performed to test for differences in the archaeal diversity based on the body location. Principal Coordinates Analysis (PCoA) based on Bray-Curtis distances was used to visualize differences between the samples from different body site. Redundancy discrimination analysis (RDA) was used to analyze the association between archaeal community composition and the body site location. RDA, alpha diversity and PCoA analysis were performed using Calypso Version 8.62 ^53^. The RSV tables obtained were used to summarize taxon abundance at different taxonomic levels. The taxonomic profiles obtained at the genus level for the samples with more than 100 reads were used to generate bar graphs for all samples.

A phylogenetic tree was constructed with the obtained archaeal RSVs from sample set 1, from the universal approach, the archaeal primer pair 519F-806R, and from the archaeal specific primer pair combination 344F-1041R/519F-806R. The alignment was performed using the SILVA SINA ^54^ and the 5 most closely related available sequences (neighbors) were downloaded together with the aligned sequences. All sequences were cropped to the same length (276 nt, from position 545 nt to 821 nt) and used to construct a tree based on maximum-likelihood algorithm using MEGA7 ^55^, using a bootstrap value of 500. The Newick output was further processed with iTOL interactive online platform ^56^.

## Declarations

### Ethics approval and consent to participate

Research involving human material was performed in accordance with the Declaration of Helsinki and was approved by the local ethics committees (the Ethics Committee at the Medical University of Graz, Graz, Austria). (Bacterial) microbiome studies of some of the samples used in this study have already been published elsewhere (oral, nose, skin samples: ^46,48,50^). Details of the ethics approvals obtained are shown there. Appendix samples and stool samples have been obtained covered by the ethics votes: 25-469 ex12/13, and 27-151 ex 14/15.

### Availability of data and material

The MiSeq amplicon sequence data was deposited in the European Nucleotide Archive under the study accession number PRJEB27023.

### Authors’ contributions

The study was designed by M.R.P. and C.M.E. G.S., H.T., V.S., E.S., B.K. and C.H. provided clinical samples. C.S. and M.R.P. prepared the 16S rRNA gene amplicons for sequencing and performed the quantitative PCR. Bioinformatics and statistical analysis were done by M.R.P. Analysis, visualization and interpretation of the data was done by M.R.P. All authors read, corrected and approved the final manuscript.

## Acknowledgements

The authors acknowledge the support of the ZMF Galaxy Team: Core Facility Computational Bioanalytics, Medical University of Graz, funded by the Austrian Federal Ministry of Science, Research and Economy (BMWFW), Hochschulraum-Strukturmittel 2016 grant as part of BioTechMed Graz. M.- R. Pausan and M. Blohs were trained within the frame of the Ph.D. program in Molecular Medicine of the Medical University of Graz.

